# DRED: A Comprehensive Database of Genes Related to Repeat Expansion Diseases

**DOI:** 10.1101/2024.09.07.611652

**Authors:** Qingqing Shi, Min Dai, Yingke Ma, Jun Liu, Xiuying Liu, Xiu-Jie Wang

## Abstract

Expansion of tandem repeats in genes often causes severe neuromuscular diseases, such as fragile X syndrome, huntington’s disease, and spinocerebellar ataxia. However, information on genes associated with repeat expansion diseases is scattered throughout the literature, systematic prediction of potential genes that may cause diseases via repeat expansion is also lacking. Here, we develop DRED, a Database of genes related to Repeat Expansion Diseases, as a manually-curated database that covers all known 61 genes related to repeat expansion diseases reported in PubMed and OMIM, along with detailed repeat information for each gene. DRED also includes 516 genes with the potential to cause diseases via repeat expansion, which were predicted basing on their repeat composition, genetic variations, genomic features, and disease associations. Various types of information on repeat expansion diseases and their corresponding genes/repeats are presented in DRED, together with links to external resources, such as NCBI and ClinVar. DRED provides user-friendly interfaces with comprehensive functions, and can serve as a central data resource for basic research and repeat expansion disease-related medical diagnosis. DRED is freely accessible at http://omicslab.genetics.ac.cn/dred, and is frequently updated to include newly reported genes related to repeat expansion diseases.

## Introduction

Repeated sequences, also known as repetitive elements, comprise more than 50% of the human genome, among which millions are short tandem repeats (STRs) with typical repeat lengths of 2–6 bp [1–3]. Although the majority of STRs are located in intergenic non-coding regions, many human coding genes also harbor STRs in exons or introns [4,5]. Copy number variation of repeat units is commonly seen among STRs, which may be caused by polymerase slippage during the DNA replication, repair, and recombination processes [6–8]. Abnormal expansion of STRs can lead to gene dysfunction at the RNA or protein level, and result in more than 40 severe inherited diseases [2,3,9–12]. Notably, RNAs with expanded repeats can independently promote phase separation and gelation, forming RNA foci in the nuclei [13–15]. Most repeat expansion related disorders are neurological, neuromuscular, or neurodegenerative diseases, such as the (CGG)_n_ repeats in fragile X syndrome, (CAG)_n_ repeats in Huntington’s disease, and (GAA)_n_ repeats in Friedreich’s ataxia [16–20]. For these diseases, the expansion of STRs is usually non-toxic when the copy numbers of STRs are below certain threshold, however, along cell division, the expansion of STRs can accumulate and become pathogenic, and result in severe symptoms. The repeat expansion diseases usually have earlier onset in descendent generations, such phenomenon is known as genetic anticipation and is a hallmark of repeat expansion diseases [18,20,21].

The majority of known disease-causal repeats are trinucleotide tandem repeats, with CAG (encoding polyglutamine) and GCG (encoding polyalanine) being the most prevalent STRs within protein coding regions [22,23]. Multiple factors at the *cis*-regulation level could promote the expansion of STRs, including adjacent to CpG islands [24], mutations in adjacent CCCTC-binding factor (CTCF) binding sites [25], as well as the presence of nearby *Alu* elements [26,27] or topological associating domains (TAD) boundaries [28].

Here, we present DRED as the first database of genes related to repeat expansion diseases. DRED not only encompasses comprehensive information on known causal genes for repeat expansion diseases, but also provides a list of predicted genes with the potential to cause diseases via repeat expansion, therefore may help researchers to identify unknown repeat expansion diseases and novel disease-causal genes.

## Database contents and construction

### Database contents

DRED contains all reported 61 genes related to 62 known repeat expansion diseases or disease subtypes collected in the PubMed or OMIM databases (**Figure 1A** and B). For each disease or disease subtype, its phenotype and general information, pathogenic gene, pathogenic repeat, repeat conservation status, pathogeny, and related references are included. Links to external data resources, such as KEGG [29], Gene Ontology [30,31], and ClinVar [32], are also provided. The expandable STRs of these 61 genes can be classified into 22 types, with CAG, CGG, GCG, and TTTCA/TTTTA as the most commonly seen expandable STR types (**Table 1**). The distributions of known disease-causal STRs are comparable across 5’ untranslated regions (UTRs), introns, and coding sequences (CDS), but are under presented in exons or 3’ UTRs of non-coding genes (Table 1). Among the known repeat expansion diseases, only spinocerebellar ataxia type 8 and oculopharyngeal myopathy with leukoencephalopathy (OPML) are caused by STRS within noncoding genes, namely *ATXN8OS* [33] and *LOC642361/NUTM2B-AS1* [34], respectively. It is worth noting that a total of 12 subtypes of spinocerebellar ataxias are related to repeat expansion, among which 7 are caused by abnormal expansion of CAG repeats encoding polyglutamine tracts in different genes [35,36].

**Table 1.**
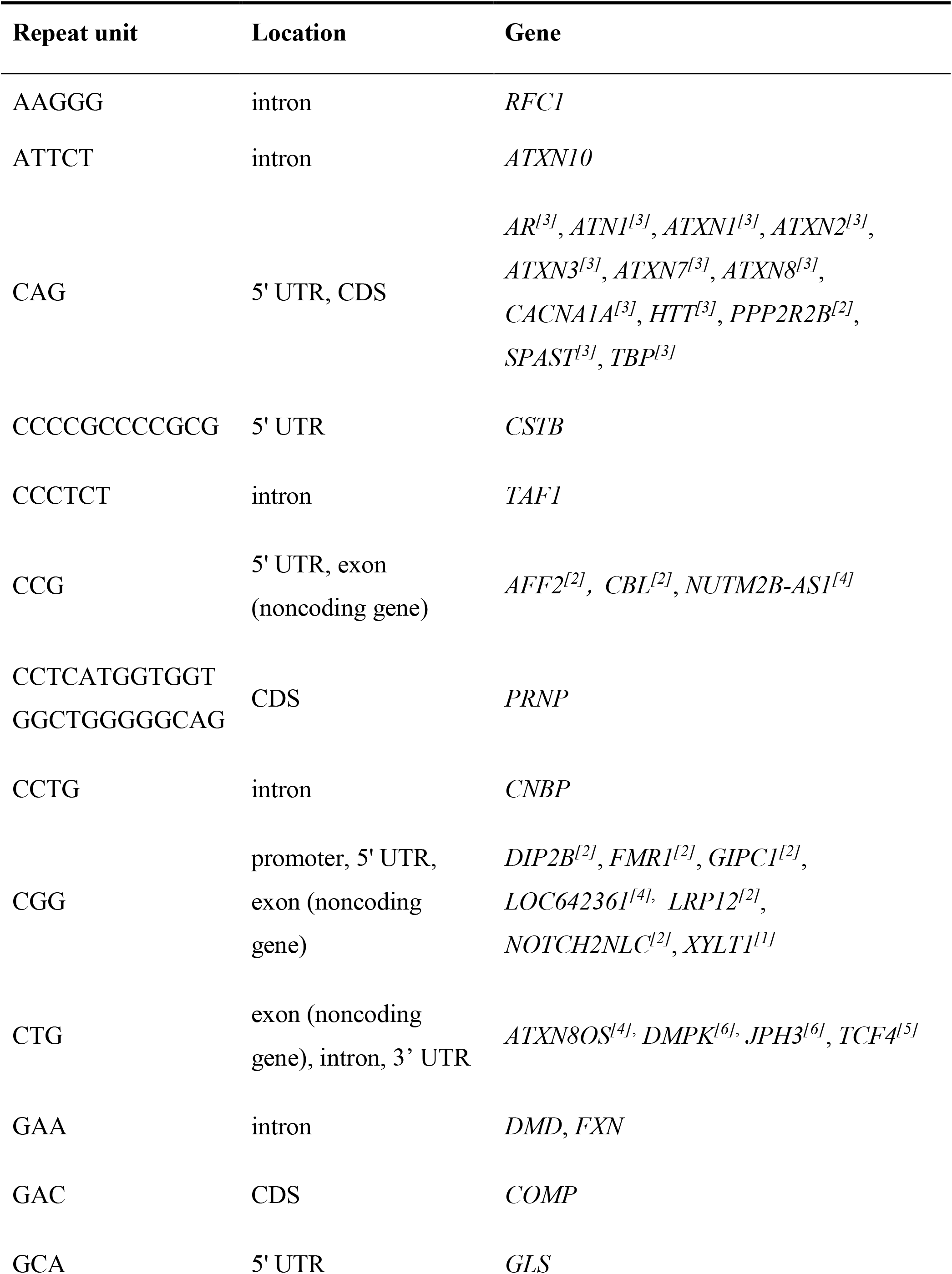
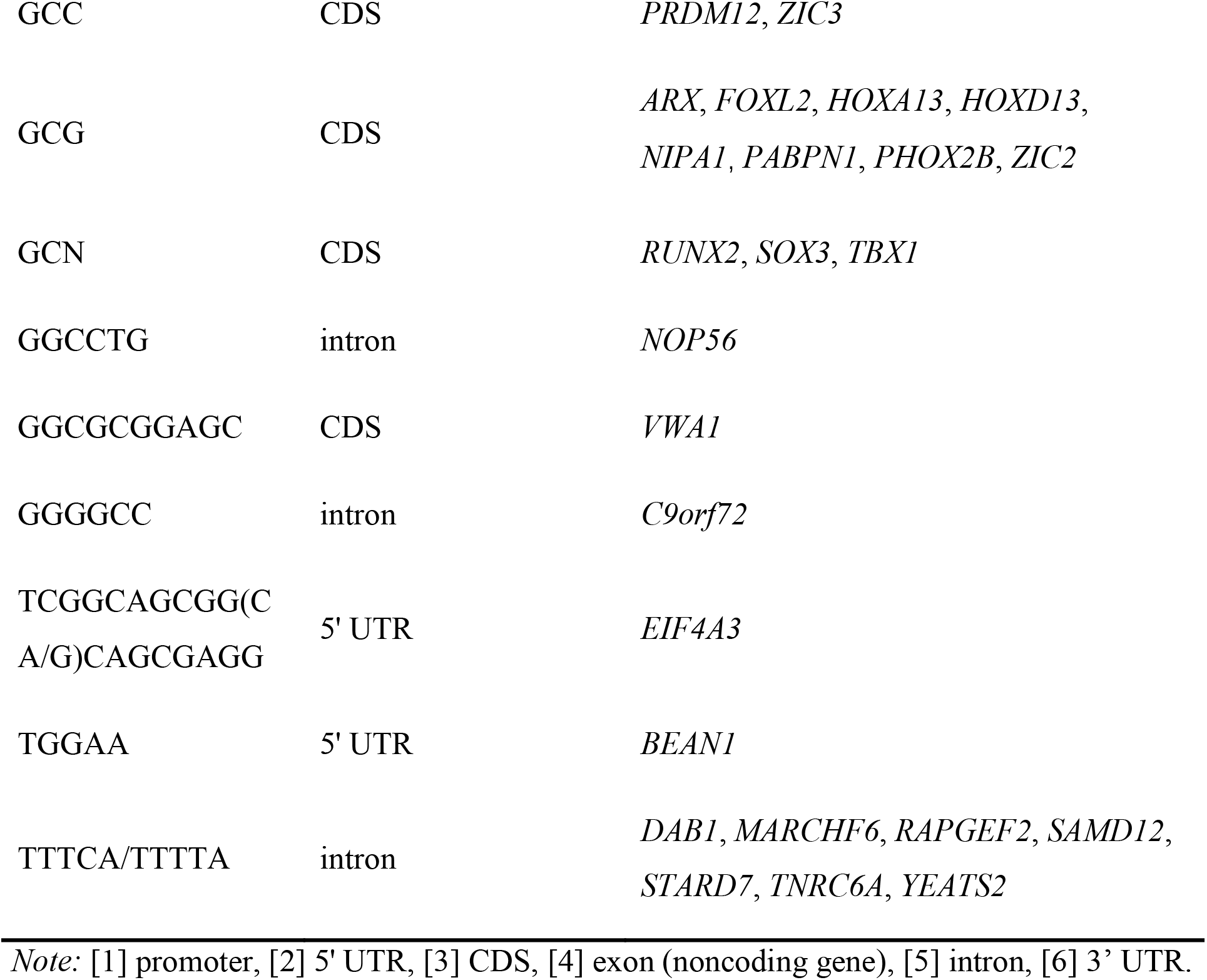
Summary of the known repeat expansion disease associated genes collected in DRED.

**Figure 1.**
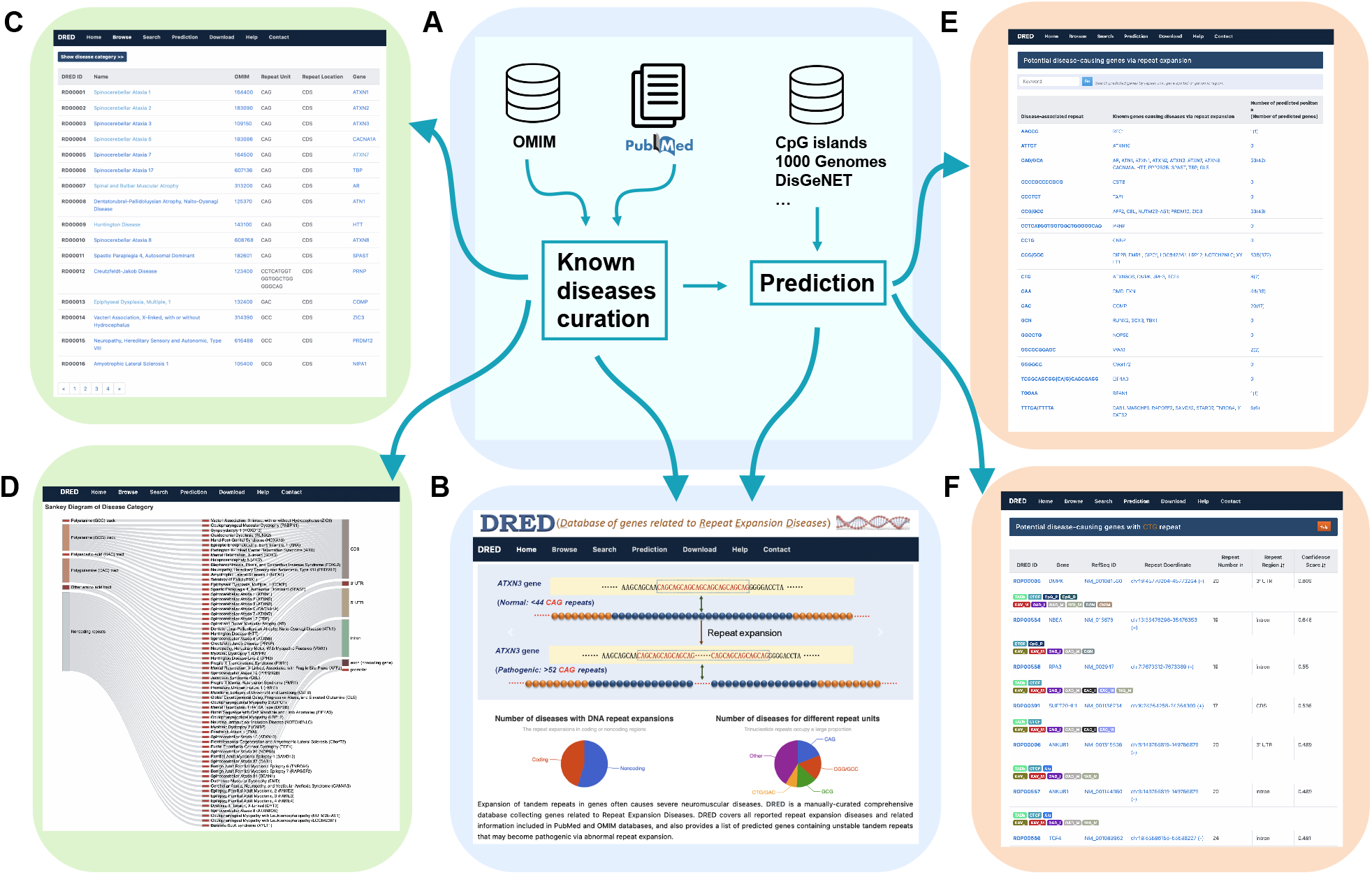
Scheme and functional illustration of DRED. **A**. Design scheme of DRED. **B**. Function overview of DRED. **C**. List of known repeat expansion diseases under the Browse function. **D**. Interactive Sankey diagram showing the categories of known repeat expansion diseases. **E**. Overview of the predicted disease-causal genes. **F**. Availability of external information for the predicted disease-causal genes.

To search for additional genes with the potential to induce diseases by repeat expansion, we collected sequence features known to contribute to the expansion of repeat sequences, and used an unsupervised machine learning algorithm to predict genes with the potential to induce diseases via repeat expansion. The features used for gene selection include: the presence of known disease-causal STRs, co-localization with *Alu* elements, CpG islands, CTCF binding sites and TAD boundaries, as well as sequence variations among populations and reported disease associations. A total of 516 candidate genes that may cause diseases via STR expansion were identified. These genes were classified by repeat types and included in the prediction section of DRED. For each predicted repeat-causal STRs, DRED provides the information on putative expandable STRs, *cis*-elements adjacent to STRs, phylogenetic conservation of STRs, variations of STRs among populations, and associated diseases. Links to the corresponding NCBI gene webpage, expression information [37], Gene Ontology annotation [38], and the UCSC Genome Browser [39] are also provided.

### Web interface and usage

DRED provides user-friendly web interfaces with comprehensive functions as described below.

#### Browse

The browse function allows users to explore the comprehensive information of all known repeat expansion diseases (Figure 1C). The 62 known repeat expansion related diseases are grouped by the features of their causal STRs, namely “Polyalanine (GCC) track”, “Polyalanine (GCG) track”, “Polyaspartic-acid (GAC) track”, “Polyglutamine (CAG) track, “Other amino acid track”, and “Noncoding repeats” (Figure 1D). For each disease listed in the Browse page, detailed description on disease phenotype, disease-causal genes, pathogenic repeat unit, repeat length, related references and other information can be obtained by corresponding links.

#### Search

The search function features a user-friendly interface that allows users to find specific information related to a repeat expansion disease. The search engine supports free-text queries, including disease names, gene symbols, repeat units, OMIM identifiers (IDs), chromosome numbers, or any keyword related to a disease. For example, entering the word “ataxia” will retrieve 18 entries with “ataxia” in the disease names or alternative disease names. An interactive 3D word cloud is provided in the search page to inform users the known repeat expansion diseases and their related genes in the database. Users can also pull up the detailed descriptions for each disease or gene by clicking on any term within the word cloud.

#### Prediction

The prediction function provides a comprehensive list of genes with the potential to cause diseases via repeat expansion. A total of 516 repeat-containing genes (477 protein coding genes and 39 noncoding genes) belonging to 19 repeat types are included (Figure 1E). Users can retrieve all predicted genes with any repeat type by clicking on either the repeat unit link or the corresponding gene count. For each gene, the detailed description, known repeat variations, as well as links to several external databases, are available via the link under DRED ID (Figure 1F). The prediction score and co-localization information of each gene with various *cis*-elements are also included. The genomic distribution features and predicted disease-causal scores of the predicted genes are similar to those of the known causal genes for repeat expansion diseases (**Figure 2A**). Gene Ontology analysis using clusterProfiler [40] and GOSemSim [41] reveals an enrichment of terms related to neural system and limb development among the 516 potential disease-causal genes (Figure 2B), which is in concert with the neurological or neuromuscular related functions of most known repeat expansion diseases.

**Figure 2.**
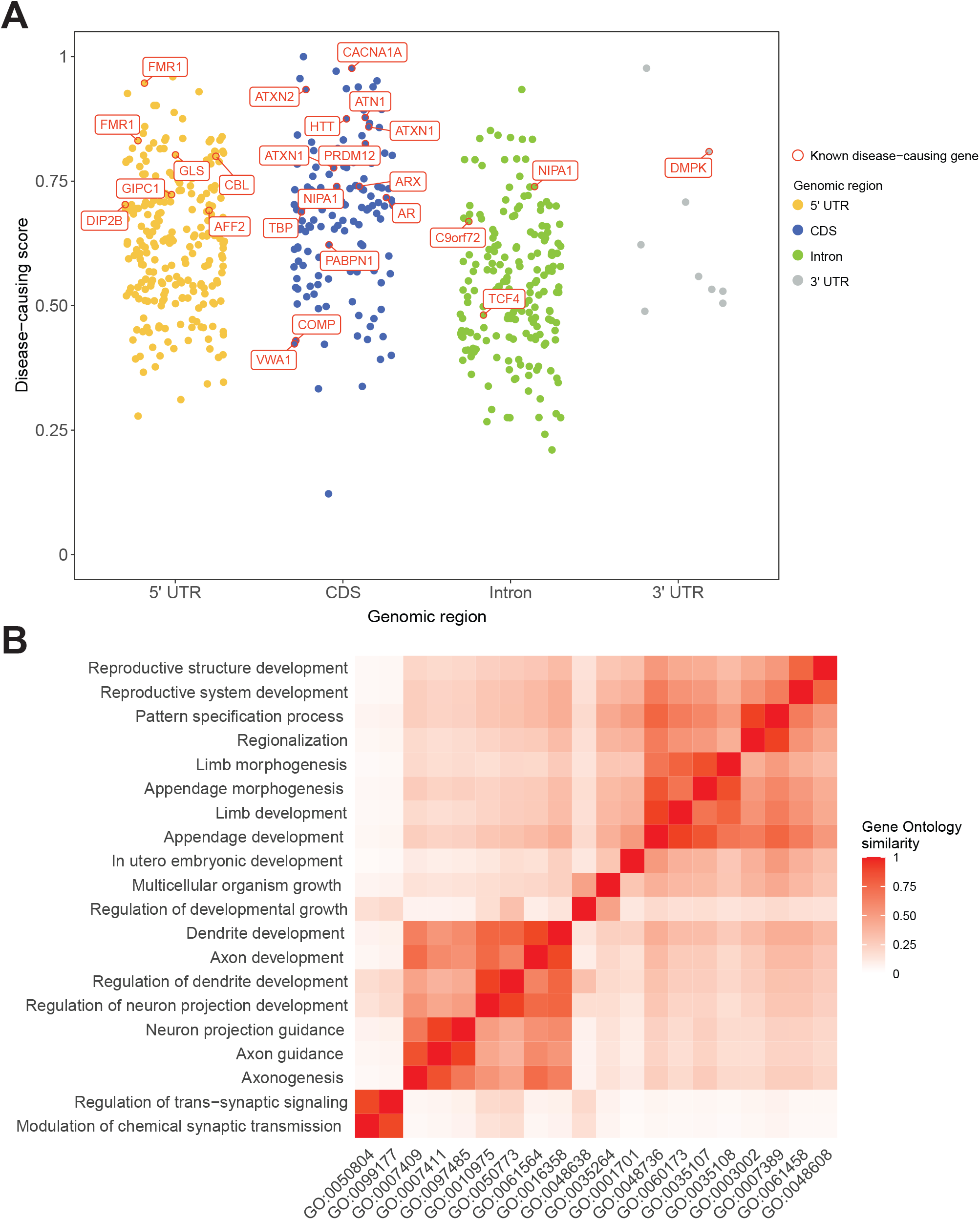
Characterization of predicted disease-causal protein-coding genes. **A**. Genomic distributions and prediction scores of the predicted disease-causal genes and the known ones. Known disease-causal genes are displayed in red boxes. **B**. Gene Ontology enrichment analysis for the 516 predicted disease-causal protein-coding genes. Shown are the top 20 enriched Biological Process terms.

#### Download

All data collected in DRED are available for local manipulation through the download function. Information on known repeat expansion diseases and predicted disease-causal genes is provided in separate downloadable files.

## Conclusion

Abnormal expansion of STRs, mainly within protein-coding genes, is the causal factor for many neurological, neuromuscular, and neurodegenerative diseases. As these diseases are inheritable and have the genetic anticipation feature across generations, early diagnosis of risky repeat carriers may help to prevent or delay the onset of the diseases, especially during the era of precision medicine. In addition, the pathogenic mechanisms of most repeat expansion diseases remain elusive, effective prevention and treatment methods are in urgent demands. Although all known disease-causal repeat expansion elements are STRs, most STRs do not have expansions or give rise to repeat expansion diseases. The current available repeat-related databases only focus on general repeat sequences in genomes, which lack comprehensive information for human repeat expansion diseases [42–45]. To meet the needs from basic research and clinical diagnosis, we developed DRED, an integrative and user-friendly database for genes related to repeat expansion diseases. DRED not only contains comprehensive information on all known causal genes for repeat expansion diseases, but also provides a list of genes with the potential to cause diseases via abnormal repeat expansion. The candidate gene list may serve as valuable resource for researchers and clinicians to identify new repeat expansion diseases and disease-causal genes, and to decipher their underlying molecular mechanisms. Continuously updated with new data every six months, DRED aims to be the premier resource for studying, diagnosing, and treating repeat expansion diseases.

## Materials and methods

### Data collection and preprocessing

#### Repeat expansion diseases in the literature

All known repeat expansion diseases were collected from the PubMed literatures [46] and OMIM [47] databases using “repeat expansion”, “trinucleotide repeat expansion”, “triplet repeat expansion”, “repeat expansion disease”, and “repeat expansion disorder” as the query words (Figure 1A). In total, 6460 publications and 14,888 disease entries (retrieved on May 26, 2023) were manually curated to remove irrelevant information. A total of 62 repeat expansion diseases with supports from PubMed publications and/or OMIM records were retained in DRED.

#### Human genetic variations

Human genetic variants in different populations of the 1000 Genomes Project [48], the Known VARiants database (Kaviar) [49], the NHLBI GO Exome Sequencing Project (ESP) [50], the sequence variation and human phenotype database (ClinVar), the Exome Aggregation Consortium (ExAC) [51,52], and the Genome Aggregation Database (gnomAD) [53] (Table S1) were collected to examine the alterations of candidate STRs among individuals, and used as a criterion for candidate disease-causal gene prediction. Picard’s LiftoverVcf (http://broadinstitute.github.io/picard/) was used to convert VCF files from the reference human genome build GRCh37 to GRCh38.

#### Alu elements, CpG islands, CTCF sites and TAD boundaries

The genomic coordinates of *Alu* elements and CpG islands were extracted according to the human GRCh38 genome assembly presented by the UCSC Table Browser (http://genome.ucsc.edu/cgi-bin/hgTables). CTCF binding peaks were obtained from 10 CTCF ChIP-seq experiments in the ENCODE project using different human tissues/cell types (Table S2) [54]. Preprocessed TAD coordinates in 40 different human tissues/cell types were downloaded from 3D Genome Browser [55], TAD boundaries of 200 kb (±100 kb centered on the boundary sites) in size were extracted using an in-house built script.

### Prediction of causal genes for repeat expansion diseases

To predict other genes with STRs that may be capable of causing diseases via repeat expansion, we firstly extracted the reported genomic and genetic features that could contribute to repeat expansion, including: (1) The presence of nearby *cis*-elements, such as *Alu* elements, CpG islands, CTCF binding sites, and TAD boundaries; (2) Variation of STR copy numbers among populations, as evaluated using the 1000 Genomes, Kaviar, ESP, ClinVar, ExAC, and gnomAD databases; (3) The implication of genes in diseases according to information in the OMIM or DisGeNET [56] databases. Details of these features are listed in Table S3. The repeat-containing genes were then selected from the GRCh38 human genome with the following criteria: (1) the genomic sequences of a gene should contain STRs with copy numbers no less than the median value of the STR copy number range associated with normal phenotypes; (2) STRs within a gene should have at least two copies of expansion in one or more records in the 1000 Genomes, Kaviar, ESP, ClinVar, ExAC, and gnomAD databases. In total, 567 STR sites from 516 genes were keep as the final prediction results in DRED.

In order to further prioritize the predicted disease-causal genes, we used Principal Component Analysis (PCA) to identify genomic features enriched among these disease-associated genes using the prcomp() function in R. In the input matrix for PCA, each row is a gene and each column is a feature, and the first principal component (PC1) captured the major variations of the input matrix (62.9%). Next, we performed a min-max normalization for genes’ coordinates on PC1 and assigned the normalized value as the disease-causal score for each gene. The corresponding weights of the 19 different features on PC1 were listed in Supplementary Table S3, and the top three weighted features were with reported STR expansion in gnomAD, the presences of CpG islands in proximity region, and overlapping of CpG islands with the repeat tracks.

### Database and web interface implementation

DRED runs on an apache web server and is implemented in PHP 5.6.31 (http://www.php.net). The server-side PHP scripts deal with SQL query for keywords submitted by users and then execute through MySQL 5.7.16 (https://www.mysql.com), and return query result via interactive web interfaces written in bootstrap 4.1.1 (https://getbootstrap.com). Interactive data visualization is supported by echarts 4.0 (http://echarts.baidu.com) and jQuery v3.3.1 (https://jquery.com). The web interface is compatible with all web browsers and may work best on Google Chrome, Firefox, or Safari.

### Gene Ontology enrichment analysis

The R package clusterProfiler v4.2.1 [40] was used to identify enriched ‘Biological Process’ Gene Ontology terms for the potential disease-causal genes. The parameters were set as follows: pvalueCutoff=0.01, qvalueCutoff = 0.01, and pAdjustMethod = “BH”. Subsequently, the semantic similarities of the top 20 enriched terms were calculated by GOSemSim v2.20.0 [41] with the following parameter: measure = ‘Wang’.

## Data availability

DRED is freely accessible at http://omicslab.genetics.ac.cn/dred.

## CRediT author statement

**Qingqing Shi:** Data curation, Software, Writing -Original Draft, Writing -Review & Editing, Visualization. **Min Dai:** Data curation, Software, Methodology, Writing -Original Draft, Writing -Review & Editing, Verification. **Yingke Ma:** Data curation. **Jun Liu:** Data curation. **Xiuying Liu:** Data curation. **Xiu-Jie Wang:** Conceptualization, Methodology, Writing -Original Draft, Writing -Review & Editing, Funding acquisition. All authors have read and approved the final manuscript.

## Acknowledgments

This work was supported by Beijing Natural Science Foundation of China (Grant No. Z200020) and the National Key R&D Program of China (Grant No. 2019YFA0802203) to XJW.

## Supplementary material

**Table S1.**
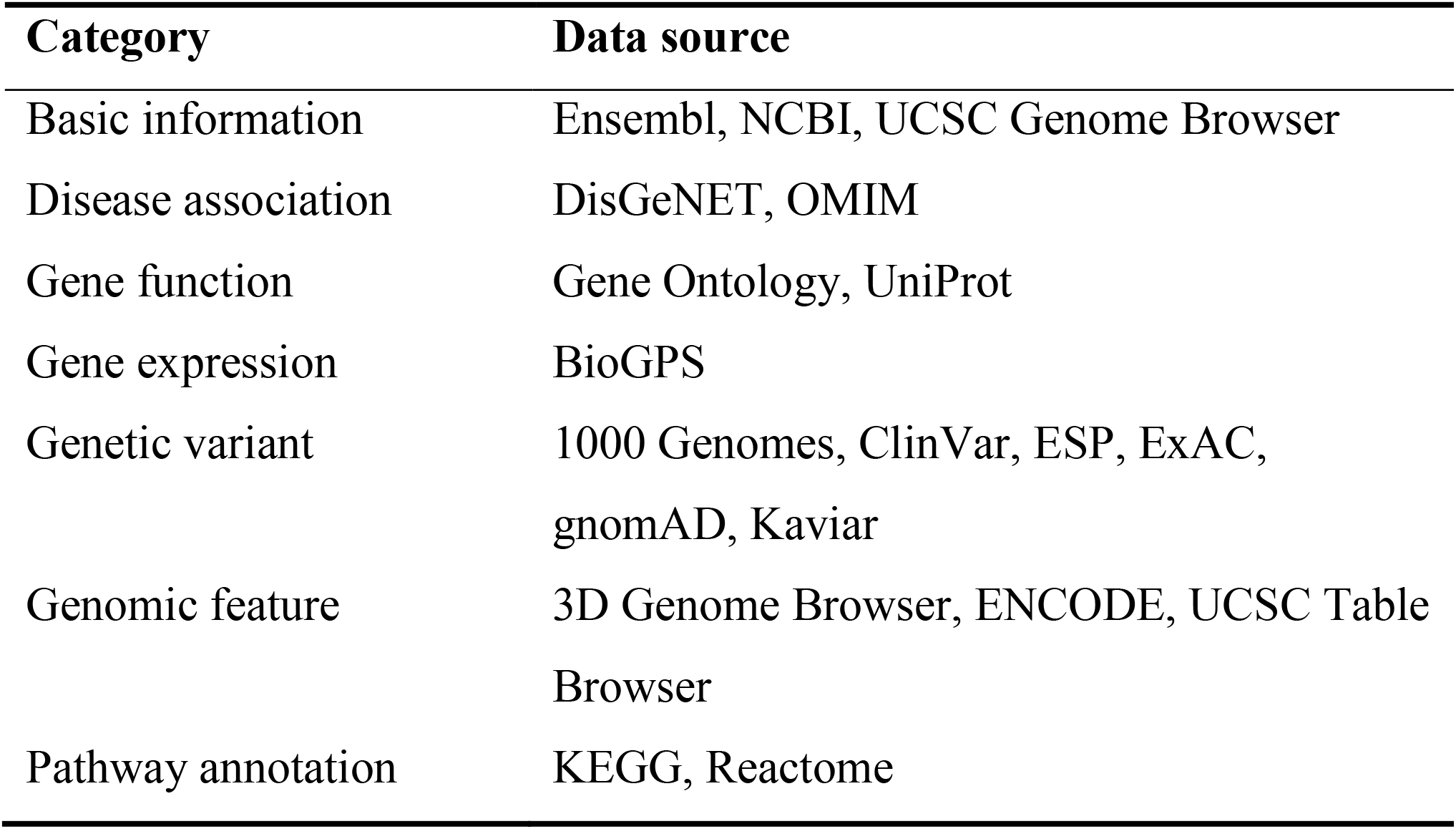
Summary of integrated data sources in DRED.

| **Category** | **Data source** |
| --- | --- |
| Basic information | Ensembl, NCBI, UCSC Genome Browser |
| Disease association | DisGeNET, OMIM |
| Gene function | Gene Ontology, UniProt |
| Gene expression | BioGPS |
| Genetic variant | 1000 Genomes, ClinVar, ESP, ExAC,<br>gnomAD, Kaviar |
| Genomic feature | 3D Genome Browser, ENCODE, UCSC Table<br>Browser |
| Pathway annotation | KEGG, Reactome |

**Table S2.** ChIP-seq data used for CTCF-binding peak identification.

| **ENCODE accession No.** | **Sample type** |
| --- | --- |
| ENCFF202NVV | Ovary from female adult (51 years old) |
| ENCFF228IAW | H1-hESCs |
| ENCFF362CYM | GM12864 cells |
| ENCFF557KXD | Neural cell originated from H1-hESC |
| ENCFF588VGU | GM12801 cells |
| ENCFF609IPZ | Human osteoblasts |
| ENCFF617JSF | HCT116 cells |
| ENCFF655TAP | Tibial nerve from female adult (51 years old) |
| ENCFF750BRK | Cardiac muscle cells |
| ENCFF788JIP | A549 cells |

**Table S3.**
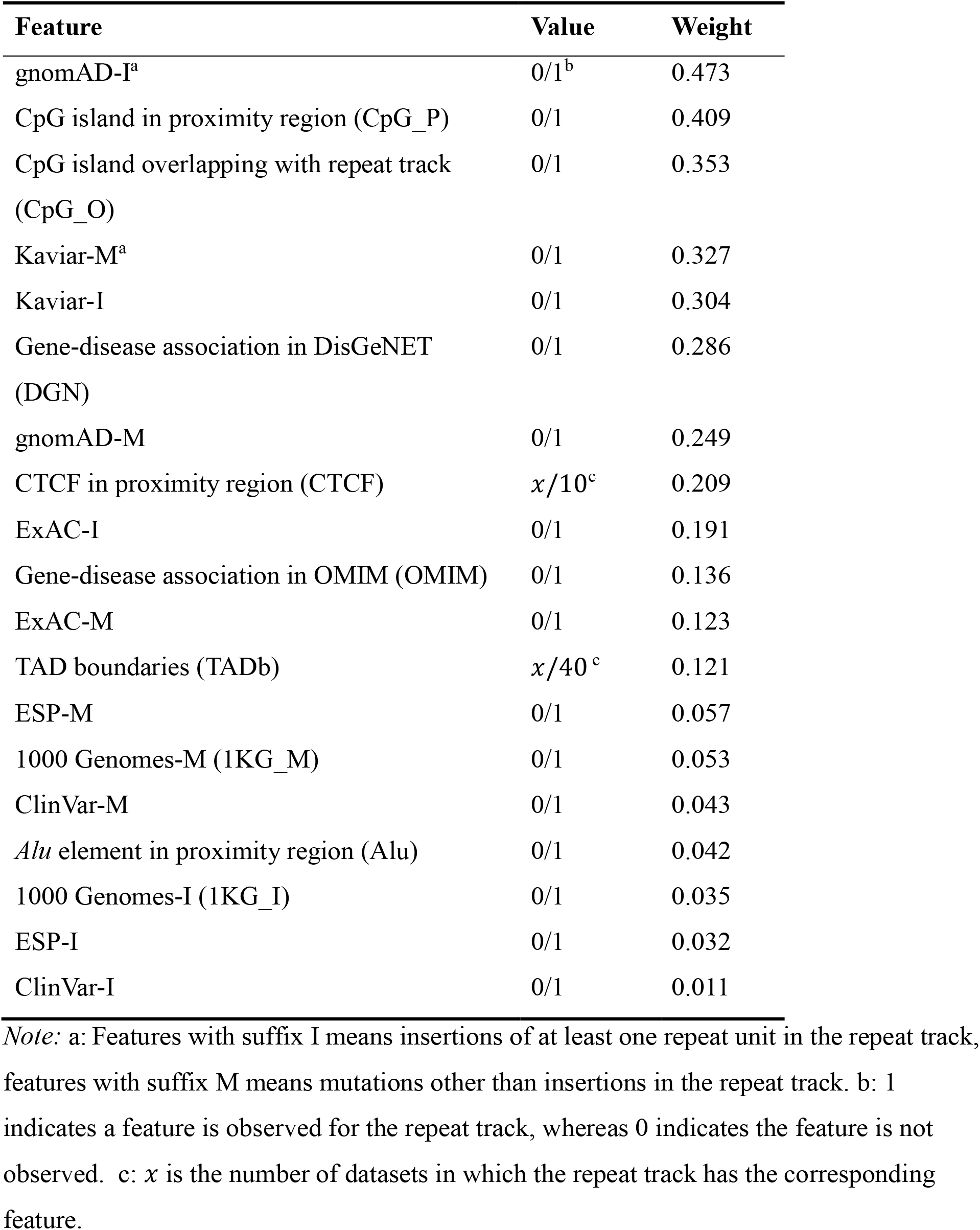
Features used for potential disease-causal gene prediction and evaluation.

| Feature | Value | Weight |
| --- | --- | --- |
| gnomAD-I<sup>a</sup> | 0/1<sup>b</sup> | 0.473 |
| CpG island in proximity region (CpG_P) | 0/1 | 0.409 |
| CpG island overlapping with repeat track (CpG_O) | 0/1 | 0.353 |
| Kaviar-M<sup>a</sup> | 0/1 | 0.327 |
| Kaviar-I | 0/1 | 0.304 |
| Gene-disease association in DisGeNET (DGN) | 0/1 | 0.286 |
| gnomAD-M | 0/1 | 0.249 |
| CTCF in proximity region (CTCF) | $x/10^c$ | 0.209 |
| ExAC-I | 0/1 | 0.191 |
| Gene-disease association in OMIM (OMIM) | 0/1 | 0.136 |
| ExAC-M | 0/1 | 0.123 |
| TAD boundaries (TADb) | $x/40^c$ | 0.121 |
| ESP-M | 0/1 | 0.057 |
| 1000 Genomes-M (1KG_M) | 0/1 | 0.053 |
| ClinVar-M | 0/1 | 0.043 |
| *Alu* element in proximity region (Alu) | 0/1 | 0.042 |
| 1000 Genomes-I (1KG_I) | 0/1 | 0.035 |
| ESP-I | 0/1 | 0.032 |
| ClinVar-I | 0/1 | 0.011 |

*Note:* a: Features with suffix I means insertions of at least one repeat unit in the repeat track, features with suffix M means mutations other than insertions in the repeat track. b: 1 indicates a feature is observed for the repeat track, whereas 0 indicates the feature is not observed. c: $x$ is the number of datasets in which the repeat track has the corresponding feature.

